# Interpretable multi-omics integration with UMAP embeddings and density-based clustering

**DOI:** 10.1101/2024.10.07.617035

**Authors:** Pol Castellano-Escuder, Derek K. Zachman, Kevin Han, Matthey D. Hirschey

## Abstract

Integrating high-dimensional cellular multi-omics data is crucial for understanding various layers of biological control. Single ‘omic methods provide important insights, but often fall short in handling the complex relationships between genes, proteins, metabolites and beyond. Here, we present a novel, non-linear, and unsupervised method called GAUDI (Group Aggregation via UMAP Data Integration) that leverages independent UMAP embeddings for the concurrent analysis of multiple data types. GAUDI uncovers non-linear relationships among different omics data better than several state-of-the-art methods. This approach not only clusters samples by their multi-omic profiles but also identifies latent factors across each omics dataset, thereby enabling interpretation of the underlying features contributing to each cluster. Consequently, GAUDI facilitates more intuitive, interpretable visualizations to identify novel insights and potential biomarkers from a wide range of experimental designs.

## Introduction

Multi-omic analyses integrate diverse data types such as genomics, proteomics, and metabolomics. Combining multiple omics modalities has the potential to uncover novel insights and biomarkers more than when each data type is analyzed alone (1, 2). The growth in high-throughput technologies has precipitated an exponential increase in omics data, underscoring the urgent need for new integration methods.

Traditional approaches to multi-omics integration have primarily focused on dimension reduction techniques. For example, methods based on Canonical Correlation Analysis (CCA) are used in RGCCA (3), while Co-Inertia Analysis is used in MCIA (4). Similarly, Bayesian Factor Analysis underpins methods such as MOFA+ (5), Negative Matrix Factorization is central to intNMF (6), Principal Components Analysis to JIVE (7), and Independent Components Analysis forms the basis of tICA (8). Although these methods have been applied across various omics datasets and biological contexts, their effectiveness and limitations vary, highlighting the need for careful consideration in their application (9).

A central limitation shared by these methods is their reliance on linear assumptions. While appropriate in some cases, this assumption can be inadequate for accurately capturing the complex, often non-linear interplay among different omics layers (10, 11). Moreover, their computational intensity poses challenges, particularly for large-scale datasets. In response to these challenges, recent advancements have shifted towards non-linear integration approaches (9, 10).

Uniform Manifold Approximation and Projection (UMAP) is a dimension reduction technique that can reveal the underlying structure in complex datasets (12). By combining manifold learning with topological data analysis, it effectively visualizes high-dimensional data in lower-dimensional spaces. UMAP stands out from other methods like PCA and t-SNE by efficiently preserving global data structures, making UMAP particularly effective in visualizing large datasets and detecting patterns that might remain hidden using other techniques (11). One of its key strengths is handling non-linear data relationships, which is crucial in multi-omics, where data types often interact in complex, non-linear ways (11). Thus, UMAP can greatly improve the interpretation and understanding of multi-omic interactions.

Despite UMAP’s strengths in addressing some of these challenges, gaps remain that require further development. For example, integrating disparate feature spaces from different omics layers can complicate the analysis and interpretation of results (13). Additionally, UMAP is often sensitive to parameters and pre-processing choices, leading to potentially misleading outputs that may not accurately reflect biology (14). The need for more robust methods that enhance UMAP’s adaptability across various data modalities and settings remains crucial, particularly for addressing the complexities of biological systems (14). Therefore, we set out to develop a model that leverages UMAP’s inherent advantages, to address the specific needs of multi-omics integration, and to benchmark its performance against six prominent joint dimension reduction multi-omics integration methods using a variety of real-world and simulated datasets.

## Results

Group Aggregation via UMAP Data Integration (GAUDI) is a method developed for the unsupervised integration of multi-omics data, capitalizing on the strengths of Uniform Manifold Approximation and Projection (UMAP) for dimensionality reduction. GAUDI begins by applying UMAP independently to each omics dataset, preserving the unique characteristics of each data type.

After processing each dataset, GAUDI concatenates the individual UMAP embeddings into a unified dataset (see Methods) and then applies a second UMAP to this concatenated dataset. This step combines the distinct omics layers into a single, lower-dimensional representation, allowing for the identification of underlying biological patterns that may not be evident in higher dimensions.

GAUDI then employs Hierarchical Density-Based Spatial Clustering of Applications with Noise (HDBSCAN) for clustering (15). Given the non-linear nature of the latent space, HDBSCAN is effective as it handles clusters of varying densities and irregular shapes without assuming a predefined number of clusters. It identifies clusters based on data density and is robust against noise and outliers, making it suitable for multi-omics datasets.

Finally, GAUDI computes metagenes using a XGBoost model to synthesize molecular features (16). This involves using XGBoost to predict UMAP embedding coordinates from molecular features (e.g., gene expressions) and identifying key features that influence the positioning of samples in the integrated latent space. Feature importance scores are then extracted using SHapley Additive exPlanations (SHAP) values to determine each feature’s contribution to the embeddings (17).

GAUDI computes metagenes for both individual omics datasets and the combined omics, integrating feature importance scores from all datasets to provide a comprehensive view of molecular features across different biological layers.

### Multi-omics integration methods comparison on artificial datasets

Next, we assessed GAUDI’s performance. In the initial phase of our validation and comparative analysis, we generated three artificial omics datasets using the *InterSIM* R package (18), following the approach described by Cantini et al. These datasets were designed with predefined reference clusters, enabling a direct comparison between the clusters generated by each integration method and the dataset’s intrinsic ‘real’ clusters. We simulated these datasets with varying cluster complexities, creating sets with five, ten, and fifteen clusters. Each cluster set was further diversified by simulating two distributions: a homogeneous distribution (‘_EQ’) and a heterogeneous distribution (‘_HET’), where the latter featured a varied number of samples per cluster.

Using these synthetic datasets, we compared GAUDI to six other leading multi-omics integration methods (9). Each method decomposed the omic matrices into factors corresponding to the artificially generated cluster counts—5, 10, and 15. Notably, intNMF and GAUDI are unique in providing default clustering outputs. For the other five methods, we derived clusters by implementing k-means consensus clustering on the factor matrices (9).

We quantified the alignment between the clusters obtained from these integration methods and the ground-truth clusters using the Jaccard index (JI), with a score of 1 indicating perfect agreement and 0 indicating complete discrepancy. Generally, all tested algorithms demonstrated strong performance across different scenarios, with most JIs exceeding 0.6 (Figure 1). intNMF, primarily designed for clustering, consistently outperformed the other five methods in almost all cases. However, GAUDI distinguished itself by achieving a JI of 1 in every scenario, irrespective of cluster count or sample distribution heterogeneity.

**Figure 1:**
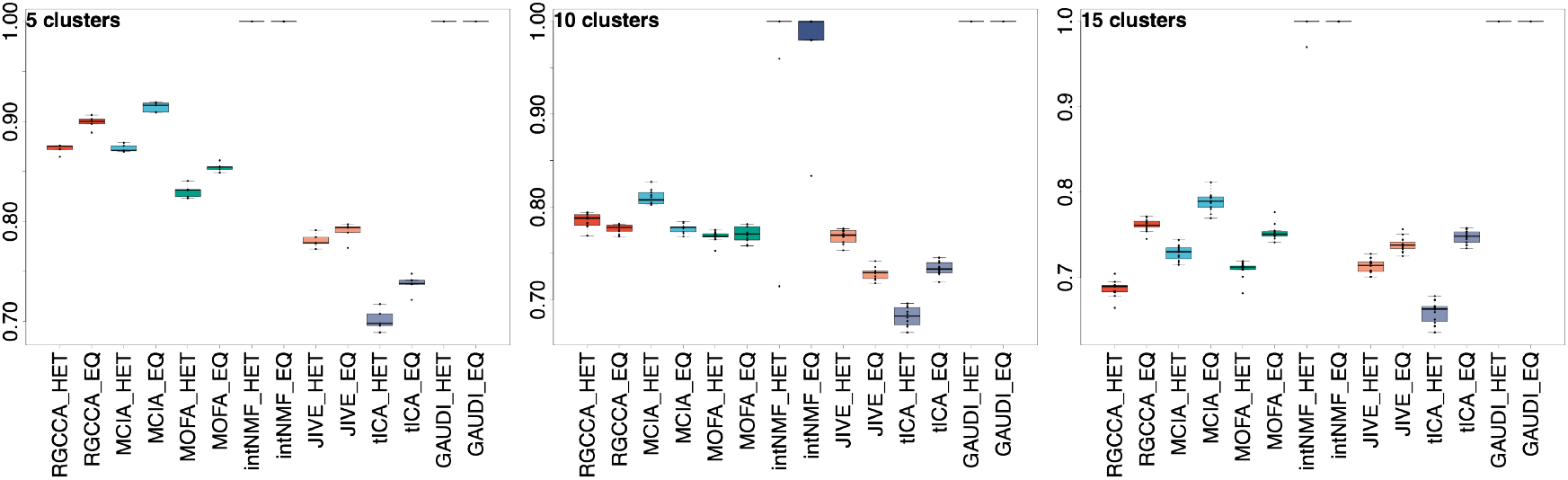
Clustering Accuracy on Simulated Multi-Omics Datasets. This figure displays boxplots illustrating the Jaccard Index (JI) scores for clusters identified by various multi-omics integration methods compared to the ground-truth clusters in the simulated data. The analysis encompasses scenarios with 5, 10, and 15 pre-defined clusters. For each method, we present results for both heterogeneous (HET) and homogeneous (EQ) cluster distributions. The analysis is based on datasets comprising 500 samples, and the depicted results are aggregated over 1,000 independent iterations of k-means clustering, ensuring robust and reliable performance evaluation.

Notably, intNMF displayed heightened variability in its performance as the number of clusters expanded and with the increase in sample size (using N=500 in our study compared to N=100 in Cantini et al.’s benchmark). In contrast, GAUDI showed remarkable consistency in clustering accuracy, unaffected by changes in cluster numbers or sample size variations (Figure 1).

These findings position GAUDI as a superior method for clustering purposes, demonstrating high accuracy and generating more condensed and differentiated clusters across all tested scenarios (Supplementary Figure 1).

### Multi-omics integration methods comparison on TCGA cancer datasets

To determine the efficacy of GAUDI to cluster multi-omics from real-world data, we used TCGA multi-omic data from eight different cancer types (19). We included gene expression, DNA methylation, and miRNA expression data, with sample sizes ranging from 170 for acute myeloid leukemia (AML) to 621 for breast cancer.

Following the benchmarking strategy outlined by Cantini et al., we decomposed the omic matrices into five latent factors and applied a Cox proportional hazards regression model to evaluate the association of these factors with survival. Consistent with previous findings, the number of factors linked to survival varied more with cancer type than with the integration method used (Figure 2A), suggesting that intrinsic biological differences across cancer types significantly influence survival outcomes (9). This underscores the importance of context-specific analysis when interpreting multi-omics data, revealing that factors such as tumor microenvironment, genetic heterogeneity, and molecular pathways play crucial roles in determining survival.

**Figure 2:**
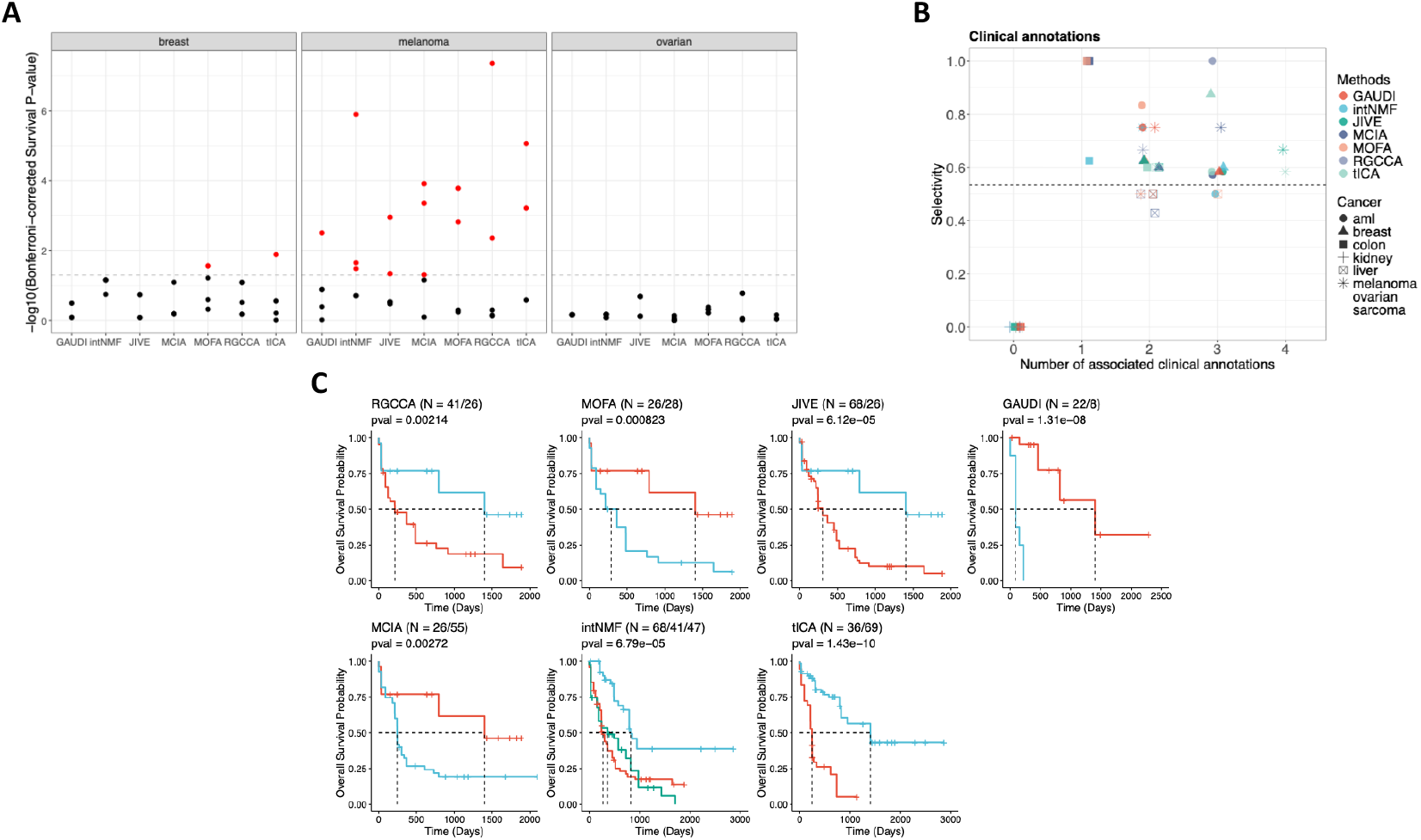
Comprehensive Analysis of Survival Prediction and Clinical Associations Using TCGA Multi-Omics Data. **A**. Bonferroni-corrected p-values for each of the 5 factors identified by multi-omics integration methods as predictive of survival, based on a Cox regression analysis. Dotted lines indicate the threshold for a corrected p-value of 0.05. Results for the remaining five cancer types are available in Supplementary Figure 2. **B**. Number of factors associated with clinical annotations in each method and the selectivity score of the factor-annotation associations. The dashed horizontal line represents the mean selectivity score. **C**. Kaplan–Meier survival curves for AML clusters with the highest and lowest median survival times identified by each method. The log-rank test is used for p-value calculations, and the sample size for each cluster is provided (N).

To extend our analysis beyond survival to clinical annotations, we tested for significant associations for the clinical annotations “age”, “days to new tumor”, “gender”, and “neo-adjuvant therapy administration” with the factors generated for each cancer type (9). The average selectivity value across all cancers and methods was 0.53 (see Methods). Methods excelling at producing clinically interpretable latent factors in each cancer type were those with the highest number of factors associated with clinical annotations and selectivity values above this average (Figure 2B).

Each method exhibited similar performance in terms of the number of generated latent factors associated with clinical annotations and selectivity scores across all cancer types, suggesting that while they identified similar numbers of associations, the relevance and impact of these associations differed depending on the specific cancer type and the underlying biological context.

To enhance the interpretability of these results and further compare the clinical applicability of each method, we examined the relationship between the clusters generated by each method and patient survival. We identified clusters with the highest and lowest median survival in each cancer type and compared their survival curves. This approach aimed to maximize survival differences based on distinct multi-omic profiles, thereby identifying molecular features predictive of extreme survival groups.

While the association between factors and survival generally depended more on cancer type than on the method used, a similar pattern emerged in the relationship between clusters and survival. Notably, none of the methods revealed significant differences in breast, colon, and ovarian cancers, but they showed comparable results for the other five cancer types (Supplementary Figure 3).

However, GAUDI stood out for its ability to detect sample groups with multi-omic profiles linked to markedly lower overall survival. This was most pronounced in acute myeloid leukemia (AML), where GAUDI identified the greatest median survival difference between the two most distant clusters (718 days, p = 1.31e-08), pinpointing a small high-risk group with a median survival of only 89 days – a threshold not reached by other methods (Figure 2C). For AML, intNMF produced clusters with identical survival medians and classified varying multi-omic profiles into similar survival groups (Figure 2C), indicating limited sensitivity at the extremes.

Using human multi-omic data, GAUDI stood out from other methods by identifying profiles associated with significantly lower overall survival times, especially in AML. These analyses demonstrate GAUDI’s enhanced sensitivity in detecting critical survival differences, which has the potential to offer valuable insights for precision medicine. In some cancers like AML, GAUDI was uniquely effective in isolating high-risk groups, an important capability that is urgently needed.

### Multi-omics integration methods comparison on single-cell datasets

Single-cell analysis has been essential for studying the biology and behavior of single cells from cell populations. As single-cell analytic techniques improve, including the ability to apply next-generation sequencing to individual cells, we have learned that cell populations previously thought to be homogeneous display considerable heterogeneity (20, 21). Thus, we tested the ability of GAUDI to detect multi-omic heterogeneity within scRNA-seq and scATAC-seq datasets. We obtained datasets containing gene expression and chromatin accessibility information across three cancer cell lines (HCT, Hela, and K562), covering 206 cells (9, 22). For this analysis, we reduced the omic datasets to two dimensions, aiming for precise classification of each cell according to its cell line.

We performed a comparative analysis including seven of the multi-omics integration methods, selected based on their C-index performance (see Methods). All methods displayed a general ability to distinguish between the three cell line types; however, MOFA+ and JIVE faced challenges in creating a clear classification in two dimensions. While the rest varied in the dispersion of their clusters, GAUDI had a misclassification rate of 1.46% (misclassifying 3 out of 206 cells), generating the most compact clusters and achieving a clear cell line identification (Figure 3A).

**Figure 3:**
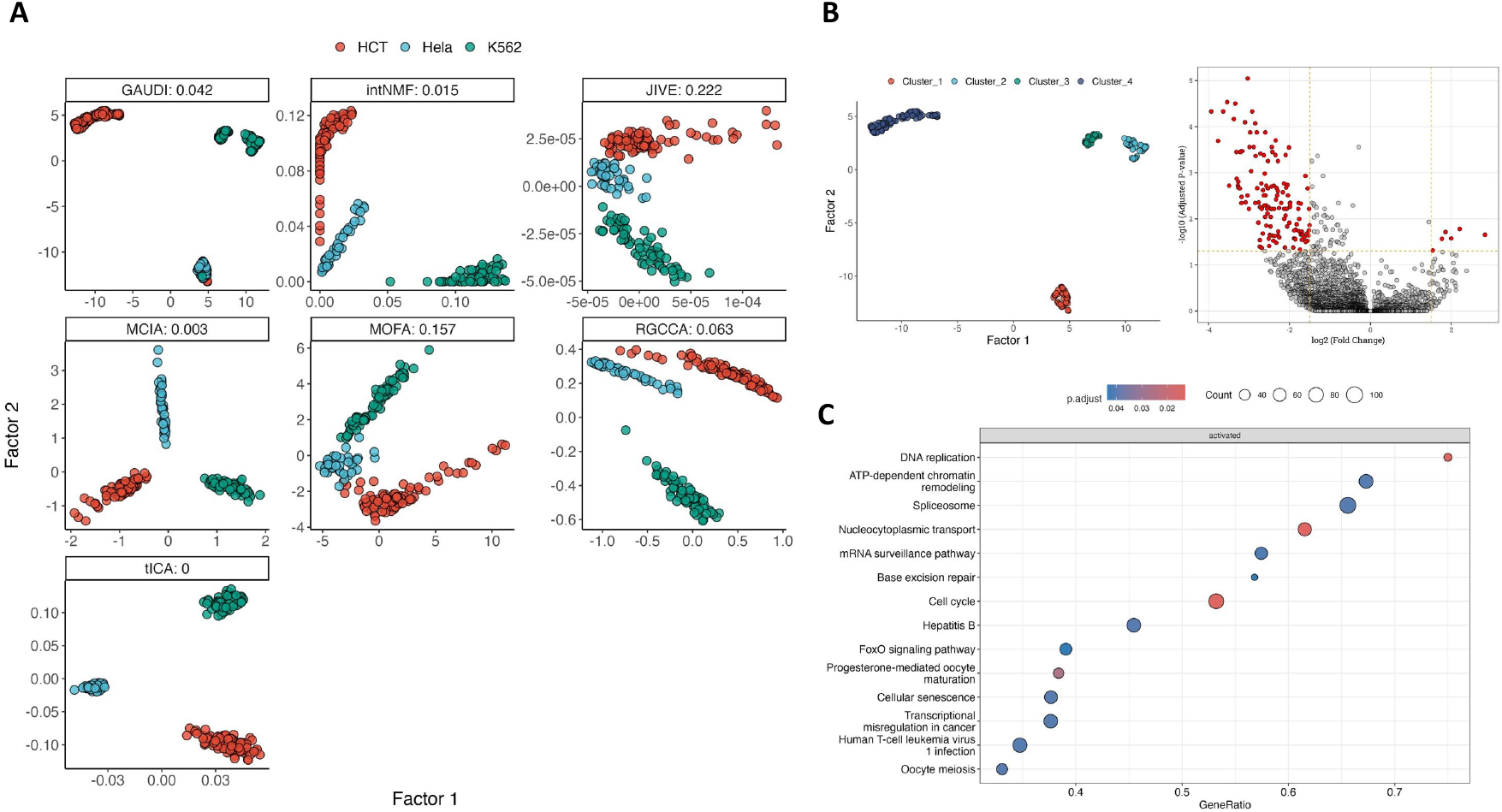
Multi-Omics Integration and Sub-structure Analysis in Single-Cell Cancer Data. **A**. Scatter plots of the two factors for each integration method (along with its C-index), color-coded by the origin of cancer cell lines. **B**. Density-based clusters by GAUDI reveal underlying sub-structures within the K562 cell line. Adjacent is a volcano plot detailing differentially expressed genes between clusters 2 and 3, highlighting genes with significant differential fold changes and FDR values. **C**. GSEA results using KEGG pathway annotations demonstrate the biological pathways differentially enriched between GAUDI-derived clusters 2 and 3 in the K562 cell line, providing insights into the functional implications of the identified substructures.

Beyond accurate cell line classification, GAUDI uniquely identified and categorized distinct multi-omic profiles within the same cell line (K562) into separate clusters (Figure 3B). We performed a differential expression analysis comparing clusters 2 and 3 within the K562 cell line, uncovering 190 differentially expressed genes with an FDR < 0.05 (Figure 3B) (23). Subsequent Gene Set Enrichment Analysis (GSEA) using KEGG pathways highlighted significant variations in DNA replication, ATP-dependent chromatin remodeling, spliceosome, and cell cycle processes between the sub-clusters, mirroring the complexity of the integrated datasets (24).

These findings underscore GAUDI’s effectiveness as a clustering tool, highlighting its ability to dissect both bulk and single-cell multi-omics datasets. GAUDI’s capacity to detect and differentiate biologically significant substructures within these datasets sets it apart from other methods that fail to capture such details.

### Multi-omics integration methods comparison on functional genomics datasets

In the final benchmark of our study, we applied the multi-omics integration methods to the Cancer Dependency Map (DepMap) Project datasets, which include a wide array of cancer cell lines characterized by multiple omic layers. This benchmark aimed to evaluate the ability of each method to integrate and interpret complex, large-scale multi-omics data with a potential impact on cancer research and therapeutic targeting.

For consistency and comparability with the TCGA benchmark (Figure 2), the DepMap evaluation focused on 258 cell lines, representing seven of the eight cancer types evaluated in the TCGA benchmark. These include AML, breast, colorectal, kidney, liver, ovarian, and melanoma lineages. We integrated four distinct omics layers—gene expression, DNA methylation, miRNA expression, and metabolomics—reducing this multifaceted data into two latent factors to test precise lineage classification for each cell.

The performance of each method was gauged by the lineage accuracy of the resulting clusters. For methods not intrinsically producing clusters, we employed k-means consensus clustering on the factor matrices. We then compared each cell line cluster against its known lineage, computing a composite score that combined the Adjusted Rand Index (ARI), which measures the similarity between the true classification and the clustering outcome, and cluster purity, an indicator of the homogeneity of the clusters (see Methods). The composite score ranged from 0 to 1, with 1 representing perfect ARI and purity, indicative of optimal clustering.

GAUDI excelled in this comprehensive test, achieving the highest composite score of 0.641, indicating a superior performance in creating condensed and pure clusters (Figure 4A). This score surpassed that of MOFA+, which followed at 0.575, showing approximately a 10% lower performance compared to GAUDI (Figure 4B).

**Figure 4:**
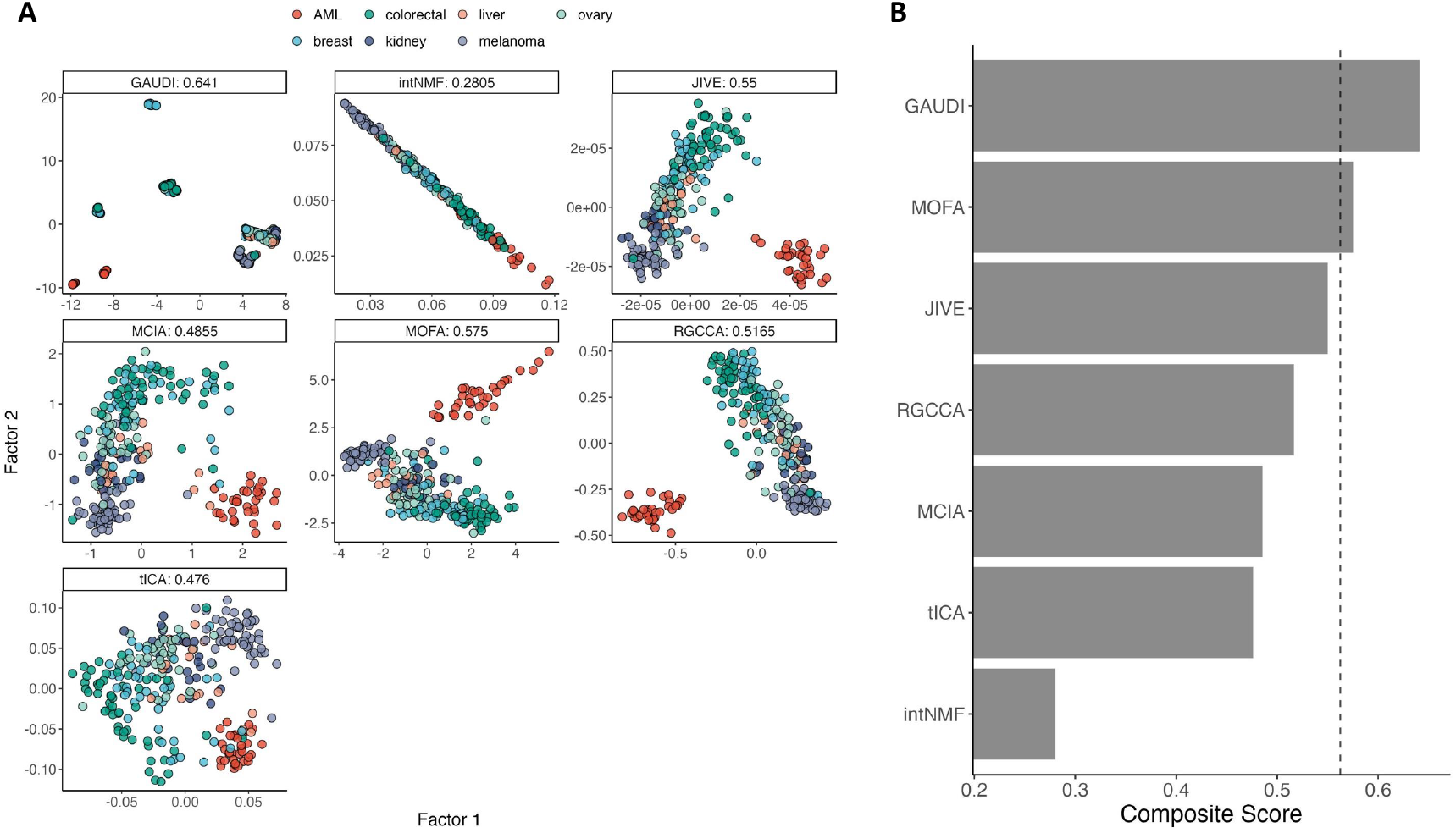
Lineage Discrimination Analysis in DepMap Multi-Omics Data. **A**. Scatterplots of the two factors derived from multi-omics integration across each method. The plots are annotated with composite scores, calculated as the average of Adjusted Rand Index (ARI) and cluster purity, which collectively assess the accuracy and homogeneity of the lineage-specific clustering. A higher composite score indicates superior performance in accurately classifying cell lineages based on the integrated multi-omics data. The cells are color-coded by its lineage. **B**. Composite scores are presented across the methods, with the vertical dashed line indicating the third quantile (75%).

The functional genomics benchmark demonstrates GAUDI’s strengths in multi-omics data integration. It demonstrated the method’s precision and efficiency, even as the complexity of the data increased.

## Discussion

In this study, we developed a novel approach to multi-omics integration and rigorously evaluated it against six leading methods benchmarked as the best joint dimension reduction methods for multi-omics integration (9). This comprehensive analysis has not only positioned GAUDI as a useful tool for multi-omics integration methods but also demonstrated that, in some instances, GAUDI performs superior to existing techniques, showcasing its proficiency in integrating complex biological data. These advancements are particularly notable in identifying survival outcomes and potential biomarkers, highlighting GAUDI’s potential to contribute to the advancement of precision medicine.

The initial benchmark using simulated datasets illustrated GAUDI’s superior capability in clustering accuracy. It achieved perfect Jaccard index scores across various scenarios, demonstrating its robustness in the face of both homogeneous and heterogeneous cluster distributions.

In the TCGA benchmark, GAUDI consistently detected sample groups with multi-omic profiles associated with markedly lower overall survival times. Notably, in acute myeloid leukemia, GAUDI identified high-risk patient groups that other methods failed to distinguish, thereby highlighting its utility in clinical prognostication. It is important to note that in this benchmark, the methods used to test the significance of clinical annotations were linear, while GAUDI’s factors are non-linear, unlike the other methods. Given this, we would expect poorer performance from GAUDI in this analysis. However, not only did GAUDI perform comparably to other methods, but it also achieved the best results in sarcoma, with a selectivity of 0.83 and two associations. These results highlight GAUDI’s capabilities not just in predicting survival but also in its association with a range of clinical annotations.

Our single-cell analysis benchmark further reinforced GAUDI’s strength. It was the only method that could detect significant sub-structures within the same cell line types, which was substantiated by differential gene expression analysis. This capability to discern finer biological details holds promise for advancing our understanding of cellular heterogeneity in cancer and other complex diseases. Additionally, these findings indicate that as the capability to analyze multi-omics in single-cell data advances, GAUDI is inherently equipped to precisely classify and manage such data. This capability offers an enhanced level of detail and interpretation, enriching our understanding of complex biological interactions at the single-cell level.

The functional genomics benchmark was perhaps the most challenging, given the scale and diversity of the datasets. Nonetheless, GAUDI demonstrated strong performance, showcasing its efficiency and accuracy even as the dimensionality of the integration increased. The method’s ability to generate precise and biologically informative clusters, even with the addition of more ‘omic layers, emphasizes its applicability to current high-throughput multi-omics datasets in cancer biology.

One limitation that emerged across the benchmarks is the computational intensity inherent in processing large-scale multi-omics data. Although GAUDI is relatively efficient, further advancements in computational strategies are needed to enhance its scalability and speed.

Additionally, while GAUDI showed promising performance on synthetic data, this success does not always translate seamlessly to real-world datasets, which tend to exhibit more noise and variability. Another challenge lies in the nature of clustering outcomes. Clustering cell types with high contrast, such as those across three different single-cell datasets, can be more straightforward than distinguishing biologically similar cells, where differences are more nuanced. This subtlety affects the clustering’s applicability and raises the question of when clustering or classification is the appropriate tool. In some research scenarios, clustering may not yield useful outcomes, making alternative methods, such as network analysis or pathway enrichment, more suitable for uncovering complex biological interactions.

Moreover, while GAUDI effectively clusters complex biological data, there are inherent limitations concerning overfitting. This concern persists even in unsupervised learning, as models might capture noise or dataset-specific patterns instead of underlying general structures. This issue is particularly pronounced in high-dimensional, noisy multi-omics data, where distinguishing the true signal from noise is challenging. Techniques such as parameter optimization, regularization, and external validation can mitigate overfitting, but they cannot eliminate this risk entirely. Therefore, we recommend users employ these strategies to enhance the robustness of their clustering results. Overfitting concerns are common across all machine learning methods, emphasizing the need for careful validation to ensure models generalize beyond the training data.

GAUDI’s performance highlights the importance of developing integration methods that not only provide statistical robustness but also maintain biological interpretability. By achieving accuracy in clustering and showing a strong association with clinical outcomes, GAUDI has proven to be a useful tool for researchers and clinicians, creating new opportunities for discovery in precision medicine.

Future work should focus on extending GAUDI’s application to other complex diseases and exploring its potential to guide therapeutic decision-making. Integrating GAUDI with other machine learning models may also improve its classification power for patient stratification and treatment response, offering a more comprehensive framework for interpreting complex datasets.

In conclusion, this study emphasizes the need for advanced multi-omics integration tools like GAUDI, which combines the strengths of UMAP with methodological improvements, to keep pace with the evolving landscape of biomedical data. Our benchmarking supports the adoption of GAUDI in future multi-omics studies and demonstrates its advantages over existing methods. Its ability to shed light on complex biological phenomena enhances our understanding of diseases and informs the development of precision therapies, helping to drive progress in biomedical research.

## Acknowledgments

We would like to thank the entire Hirschey lab for thoughtful feedback. We would also like to acknowledge funding support from the NIAID 1R38AI40297 (DKZ), 1K38CA282963 (DKZ), and from developmental funds of the Duke Cancer Institute as part of the Cancer Center Support Grant P30 CA014236 (MDH), from the National Institutes of Health and the NIA R01AG045351 (MDH) and 1R21AG080334 (MDH).

## Methods

### UMAP embedding integration for optimizing GAUDI

To optimize the UMAP embedding integration strategy for our GAUDI method, we systematically assessed various approaches to improve data segregation into biologically relevant clusters. This analysis was performed in Python using a custom script, “umap_integration_methods.py”, which is available in our source code repository for reproducibility.

For the benchmark, we used gene expression and miRNA expression datasets from DepMap, leveraging cell line lineage information to evaluate the resulting clusters. UMAP was first applied to each omics dataset to generate individual embeddings. We then explored several integration techniques, including intersection, union, subtraction, and concatenation of these embeddings. Additionally, we performed UMAP on the concatenated raw omics datasets and applied joint matrix factorization to the UMAP embeddings, testing the integration techniques on both the concatenated UMAP and the shared matrix embeddings.

Each integration (a total of 10) was clustered using HDBSCAN, and the effectiveness was evaluated based on cluster purity and silhouette scores (Supplementary Figure 4). This process identified the concatenation of individual UMAP embeddings followed by a combined UMAP as the most effective technique for our GAUDI method.

### Multi-omics data simulation

The multi-omics datasets in our study were simulated using the *InterSIM* package, available on CRAN (18). This package creates interrelated datasets that emulate the complexity of real-world data, based on DNA methylation, mRNA gene expression, and protein expression from TCGA ovarian cancer data. For our analysis, we generated 500 simulated samples, increasing the sample size of 100 used in Cantini et al.’s paper. As described in Cantini et al. benchmarking workflow, we created five, ten, and fifteen clusters of equal and randomly varying sizes.

### Latent space clustering

For those benchmarked methods that do not inherently generate clusters (excluding intNMF and GAUDI), we employed k-means clustering on the latent space matrix (i.e., factor matrix), aligning with the approach outlined by Cantini et al. Given the stochastic nature of k-means clustering, we executed the clustering process 1000 times. Subsequently, we derived a consensus clustering by determining the most common sample-to-cluster associations across these iterations.

### Clustering quality metrics

The Jaccard Index is a statistic used for gauging the similarity and diversity of sample sets. The index measures the similarity between sample sets by dividing the number of observations common to both sets by the total number of distinct observations in both sets. Its value ranges from 0 to 1, where 0 signifies no similarity and 1 denotes that the sets are identical.

The C-index is a clustering evaluation metric that assesses the goodness of a clustering result in relation to the within-cluster variation and the separation between clusters. A lower C-index value indicates a better clustering structure, i.e., small intra-cluster distances and large inter-cluster distances.

The Adjusted Rand Index (ARI) quantifies the congruence between two clustering solutions, taking into account the possibility of chance groupings. It accounts for both true positives and false positives within the clustering process. An ARI of 1 signifies an impeccable correspondence between the two clustering solutions, whereas an ARI of 0 indicates that the clustering is no better than a random assignment.

Cluster purity is a simple and transparent evaluation measure. It assesses the homogeneity of clusters with respect to the class distribution (e.g., lineage). The purity of a cluster is achieved when most of its members belong to the same class, with a purity score ranging from 0 (no purity) to 1 (complete purity).

To evaluate the clusters in the DepMap benchmark, we defined composite scores as the average of ARI and cluster purity, collectively assessing the accuracy and homogeneity of the lineage-specific clusters.

Finally, the silhouette score is used to measure how similar a sample is to its own cluster compared to other clusters. The silhouette score for a sample is calculated based on how much closer it is to members of its own cluster versus members of the nearest cluster to which it does not belong. The goal is to have a high silhouette score (with 1 being the highest value), which indicates that the sample is well matched to its own cluster and poorly matched to neighboring clusters.

Each of these metrics contributes to a comprehensive evaluation framework, providing us with a multifaceted understanding of the clustering quality and the effectiveness of each method in grouping multi-omics data.

### Selectivity score

The selectivity score, as defined by Cantini et al., quantifies the specificity of the associations between latent factors and clinical annotations. This score achieves a maximum of 1 when each latent factor associates with a unique clinical or biological annotation, reflecting a highly specific one-to-one relationship. Conversely, the score approaches a minimum of 0 when latent factors are broadly and non-specifically associated with multiple annotations. An ideal method would aim to maximize the number of meaningful, unique factor-annotation associations while ensuring the selectivity score remains high, thereby achieving specificity without excessive generalization (9).

### Cluster gene differential expression analysis in single-cell data

To explore molecular differences between clusters 2 and 3 identified by GAUDI, within the K562 cell line, we performed a Limma differential expression analysis using the POMA R package (23, 25). Our analysis sought to detect genes exhibiting significant expression differences between the clusters, considering an adjusted p-value of 0.05 as the threshold for significance. A volcano plot was also created to illustrate these differences, with a fold-change cutoff of 1.5 and the same p-value threshold to emphasize the most significant changes.

After this, we conducted an enrichment analysis using the R package clusterProfiler to understand the biological implications of the differentially expressed genes, focusing on KEGG pathways (26). This analysis, employing 10,000 permutations and an FDR-adjusted p-value of 0.05, helped to contextualize the gene expression patterns within broader biological pathways.

For an in-depth understanding of the benchmarking framework employed in our study, we recommend consulting the detailed exposition provided in the paper by Cantini et al.

## Data availability

The simulated data are produced using the R package *InterSIM* and can be reproduced using our quarto document “*01_comparisons_on_simulated_data*.*qmd*”. The cancer TCGA data were downloaded from http://acgt.cs.tau.ac.il/multi_omic_benchmark/download.html. The single-cell data are available in the *“data/”* folder of our GitHub repository (https://github.com/hirschelab/umap_multiomics_integration). Biological annotations were obtained from Cantini et al. GitHub repository and can be found in our GitHub repository as well. Finally, Cancer Dependency Map datasets were obtained from https://depmap.org/portal/download/custom.

## Code availability

The source code necessary to reproduce the analyses and figures presented in this paper is fully available to the scientific community at our GitHub repository, https://github.com/hirscheylab/umap_multiomics_integration, ensuring transparency and ease of access.

In addition to the source code, we have developed a dedicated R package for the method presented in this paper, GAUDI (https://github.com/hirscheylab/gaudi). This package is designed to facilitate the easy implementation of our method by other researchers. The package includes comprehensive documentation and examples to assist users in implementing our method in their own research.

We encourage researchers to use these resources for their work and welcome any contributions or feedback to improve the utility and functionality of this method.

