## Supplementary Material for "Interpretable multi-omics integration with UMAP embeddings and density-based clustering"

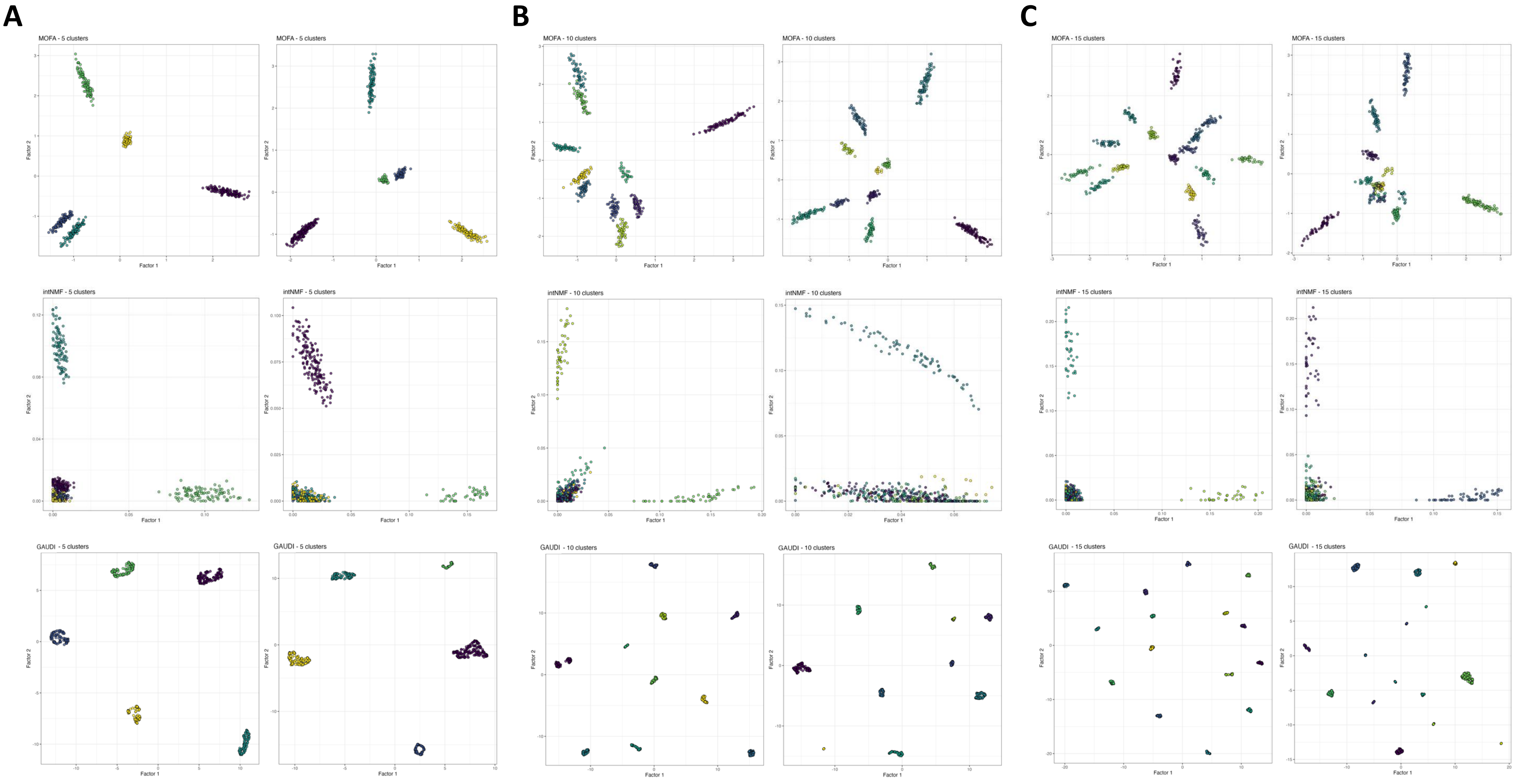


**Supplementary Figure 1: MOFA, intNMF, and GAUDI 2-dimensional Factorizations on Simulated Multi-Omics Datasets.** This figure displays scatterplots illustrating the 2-dimensional factorizations by MOFA (top), intNMF (middle), and GAUDI (bottom) in the simulated data. The analysis encompasses scenarios with 5 (**A**), 10 (**B**), and 15 (**C**) pre-defined clusters. For each method, we present results for both homogeneous (left) and heterogeneous (right) cluster distributions. The analysis is based on datasets comprising 500 samples. GAUDI shows the most condensed and differentiated clusters across all tested scenarios, achieving a Jaccard Index of 1 in every scenario, irrespective of cluster count or sample distribution heterogeneity.

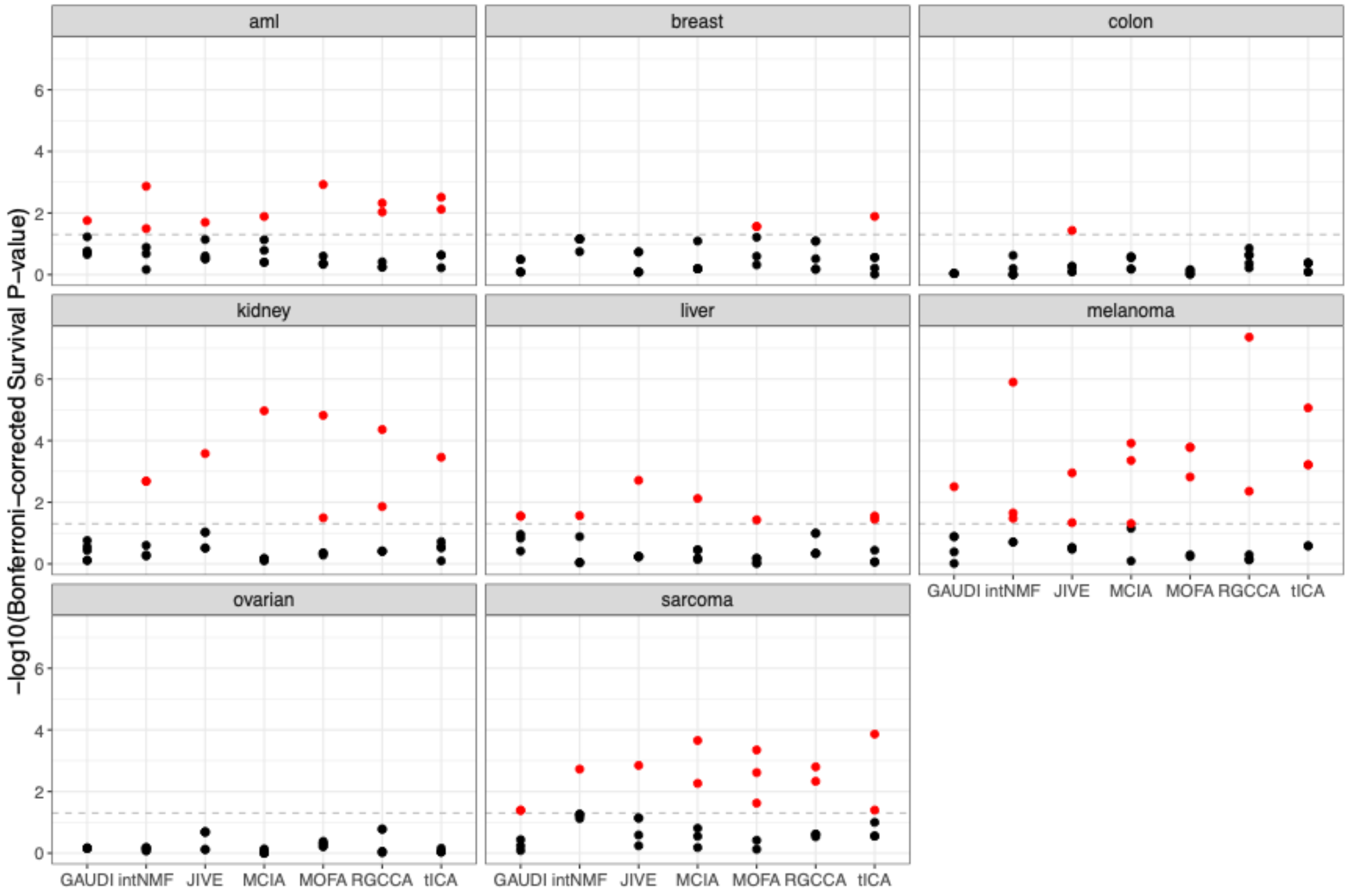


**Supplementary Figure 2: Comprehensive Analysis of Survival Prediction Using TCGA Multi-Omics Data.** Bonferroni-corrected p-values for each of the 5 factors identified by multi-omics integration methods as predictive of survival across all cancer types, based on Cox regression analysis. Dotted lines indicate the threshold for a corrected p-value of 0.05.

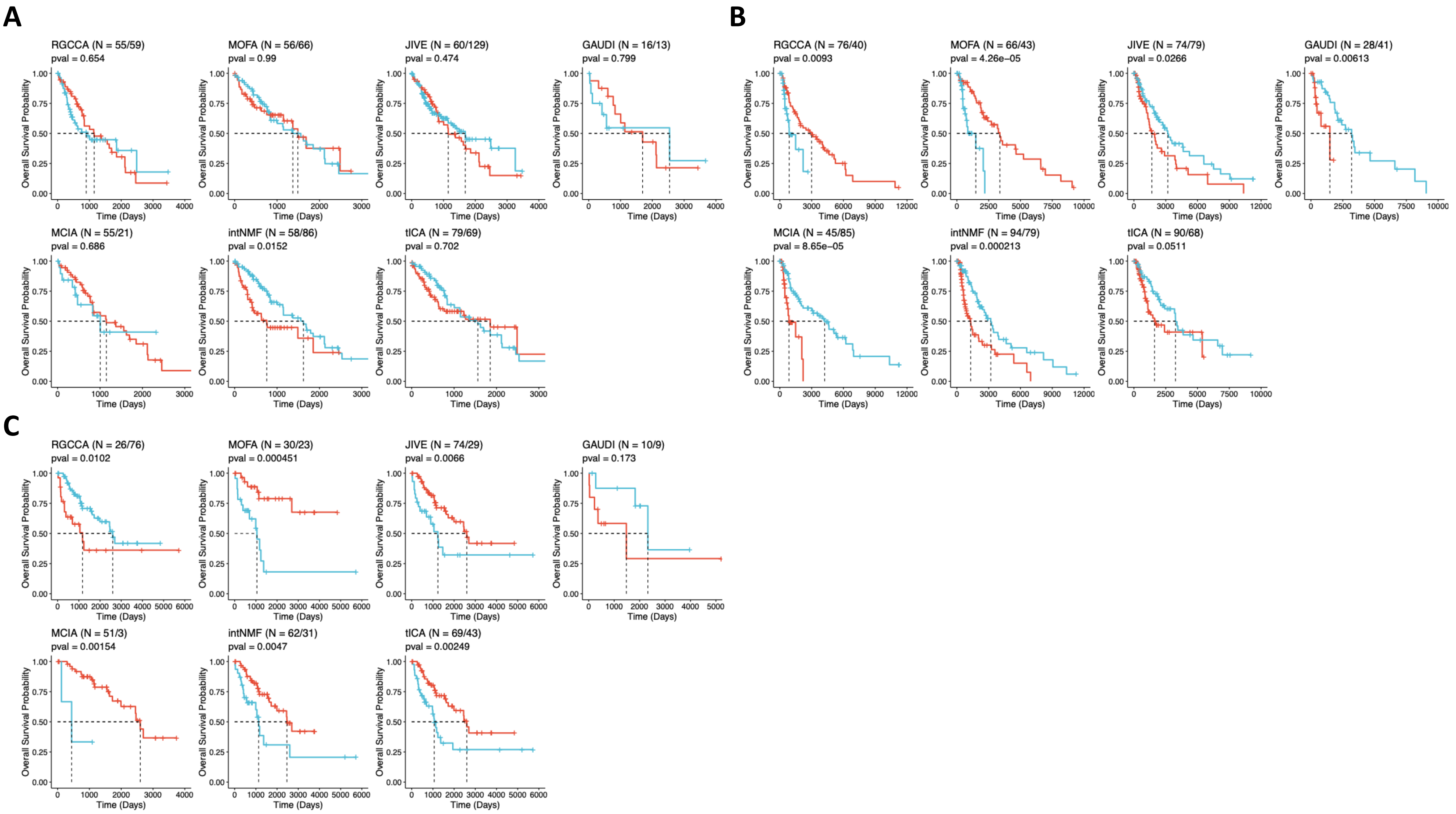


**Supplementary Figure 3: Comprehensive Analysis of Survival Clusters Using TCGA Multi-Omics Data.** Kaplan–Meier survival curves for liver (**A**), melanoma (**B**), and sarcoma (**C**) clusters identified by each method in TCGA datasets. The log-rank test is used for p-value calculations, and the sample size for each cluster is provided (N).

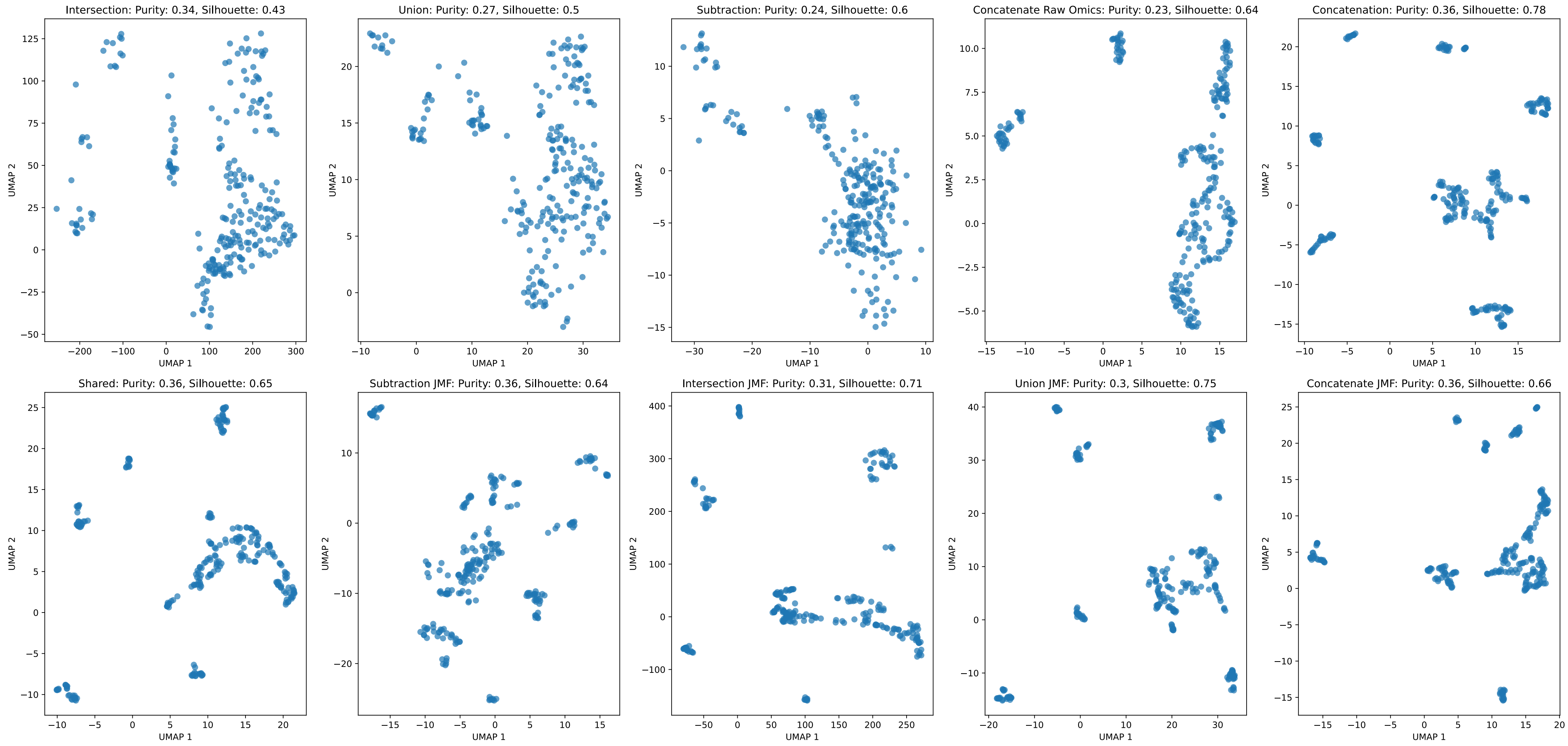


**Supplementary Figure 4: Assessment of Various UMAP Embedding Integration Techniques for Optimizing the GAUDI Method.** Each technique uniquely combines UMAP embeddings and is evaluated based on cluster purity (aligned with cell lineage) and silhouette score. The most effective method emerged as the concatenation of individual embeddings followed by a unified UMAP (top-right corner). For a comprehensive understanding, refer to the "scripts/umap_integration_methods.py" script available in our source code repository.
